# Virtual screening of inhibitors for the Zika virus proteins

**DOI:** 10.1101/060798

**Authors:** S. Feranchuk, U. Potapova, S. Belikov

## Abstract

The NS3 protease and NS5 polymerase structures of the Zika virus were constructed and processed using a virtual screening pipeline. MM-PBSA calculations show that some of the ligands found by the pipeline demonstrate a good affinity to vulnerable sites in both proteins. The anti-hypertension drug eprosartan is among the selected ligands; and inhibition of the tick-borne encephalitis virus has already been confirmed in vivo by a previous study. In the present study phytochemicals bisabolol and andrographolide are suggested as the candidates for antiviral therapy.

## Introduction

The most promising targets for inhibition of flaviviruses are polymerase and protease [Parkinson 2010]. Virus protease is a part of the NS3 protein which forms a complex with the NS2B cofactor [Falgout 1991]. The NS2B protein is needed to increase the selectivity of the NS3 protein protease domain [Aleshin 2007]. The NS2B/NS3 protease is oneof the most conservedproteins among flaviviruses [Billoir 2000].

Virtual in silico screening of ligands is a powerful tool for identifying potential drug candidates for the treatment of infections. The pipeline of virtual screening requires knowledge of the vulnerable binding sites in the target proteins. According to our previous study of the pathogenic properties of tick-borne encephalitis virus [Potapova 2012, Belikov 2014],the vulnerable site in NS3 polymerase is the interface between the NS3 protein and the NS2B cofactor; this is confirmed by other authors [Shiryaev 2011]. In the NS5 polymerase the vulnerable cavity is detected in the experiments with a site directed mutagenesis [Zou 2011] and this is confirmed by our unpublished study of NS5 protein dynamics.

The structures of proteins constructed by homology modeling and refined by molecular dynamics are reliable for virtual screening, as in the PDB there are several structures of both flavivirus proteins. Virtual screening requires a lot of computer resources, and this narrows the choice of software for protein-ligand docking and the choice for a database of ligands.Autodock Vina [Trott 2010] is a fast tool for protein-ligand docking; in our pipeline it is combined with a tool to estimate the docking quality by a statistical approach. The approved drugs and some phytochemicals were selected from adatabase of ligands, as all the ligands can be accessed easily and have additional advantages as candidates for infection treatment agents.

Reliable estimates of ligand binding energy can be obtained only by molecular dynamics simulations with a calculation of binding energy using the solution of the Poisson-Boltzmann equation on the frames of simulation (MM-PBSA method). The Amber package for molecular dynamics [Case 2005] was used for this stage of the virtual screening pipeline, as it is the most powerful tool for this kind of analysis.

## Methods

### 2.1 Models

Models of Zika virus proteins where obtained by homology modeling using the Nest program [Xiang 2006] from the following data:

NS5-sequence BAP47441.1 model pdb 4K6M

NS3-sequence BAP47441.1 model pdb 3E90

Models were processed by the standard molecular dynamics pipeline with the explicit solvent by Amber package (protonation by pdb2pqr - minimization - heating - density relaxation - production run). For both proteins, the frames of the molecular dynamics simulation after 200 ps and after 5.2 ns of the production run were selected as the two targets for virtual screening.

### 2.2 *Database of ligands*

Two databases were considered-DrugBank ligands and 2100 phytochemicals obtained by an analysis of references to molecules in Medline publications abstracts database, combined with annotation of Pubchem and structures from the Zinc database of drug-like ligands.

### 2.3 *Virtual screening*

Virtual screening was performed by Autodock Vina in a box of 20 Angstroms in size and with the center located as follows:

NS3-SER153 (site of cofactor NS2B binding)

NS5-GLU547 (sensitive pocket in the polymerase)

For each protein, two models were processed as mentioned above. After running Vina, each conformation in the screening results was processed by our unpublished algorithm which estimates the quality of the complex by statistical weight of atom combinations in the interface. The total score was averaged by two models of the receptors. The best ligands from both databases were selected for the MM-PBSA runs.

### 2.4 *MM-PBSA*

The model of the ligand in the complex was prepared by standard Amber utilities, with the use of the Mulliken empiric algorithm to assign partial charges to the atoms. The estimate of the binding energy was obtained by standard Amber scripts. The production run for 5 ns with explicit solvent was performed and 100 frames from the trajectory were selected. The binding energy was obtained by solving the Poisson-Boltzmann equation as is recommended in the Amber pipeline.

## Results

10 selected ligands for NS3 protease gave MM-PBSA energies from -21 to -10 kCal/mol 20 selected ligands for NS5 polymerase gave MM-PBSA energies from -24 to -9 kCal/mol Eprosartan (Fig. 1) was selected for demonstration as it was previously selected by virtual screening against tick-borne encephalitis virus protease and is positively tested in vivo with virus-infected mice.

Bisabolol (Fig. 2) and Andrographolide (Fig. 3) were selected for demonstration as these ligands were among the best in the results for NS3 protease and NS5 polymerase.

**Fig. 1.**
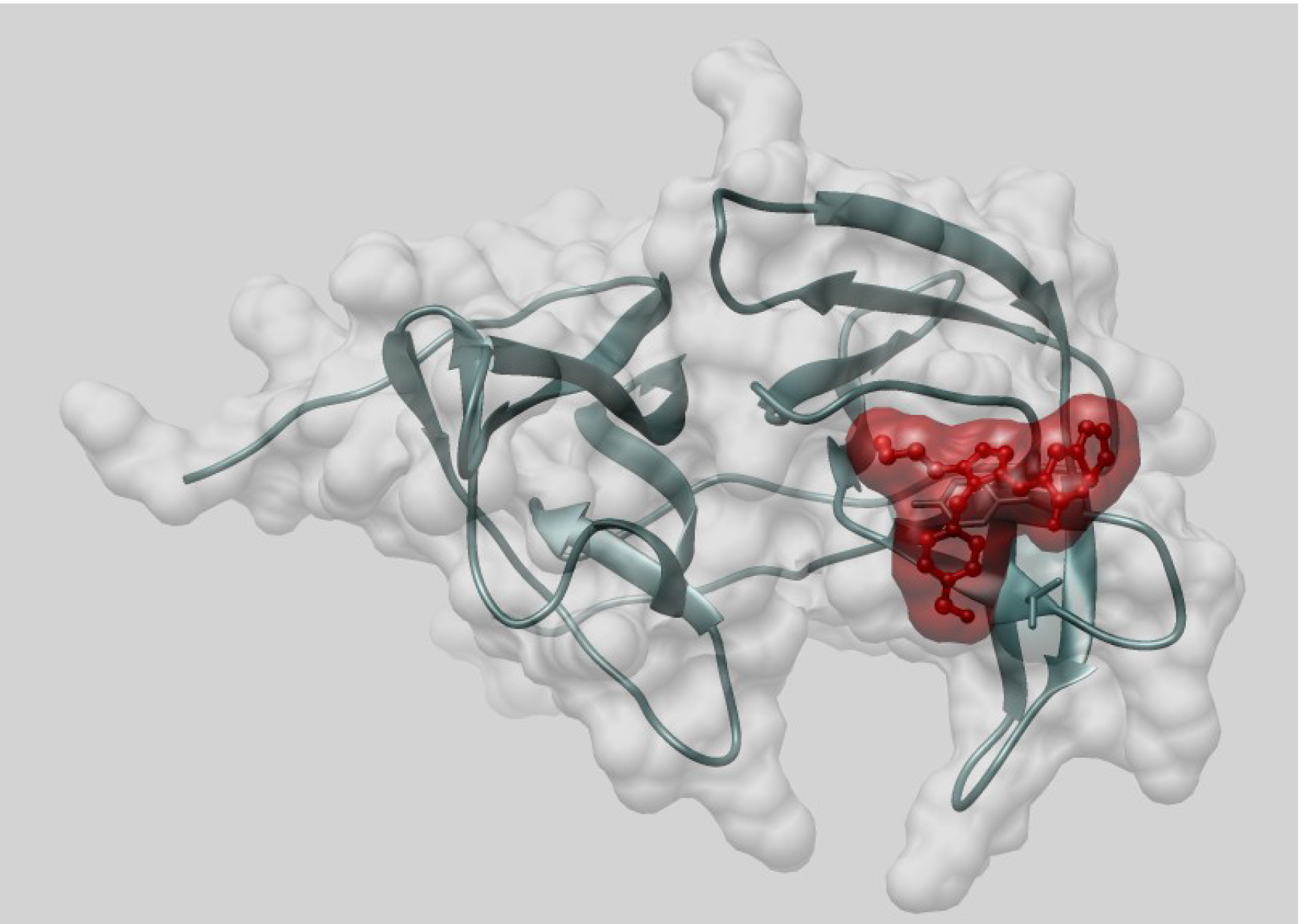
ZINC29319828 Eprosartan bound to Zika virus NS3 protease(MM-PBSA Energy -16.62 kCal/mol)

**Fig. 2.**
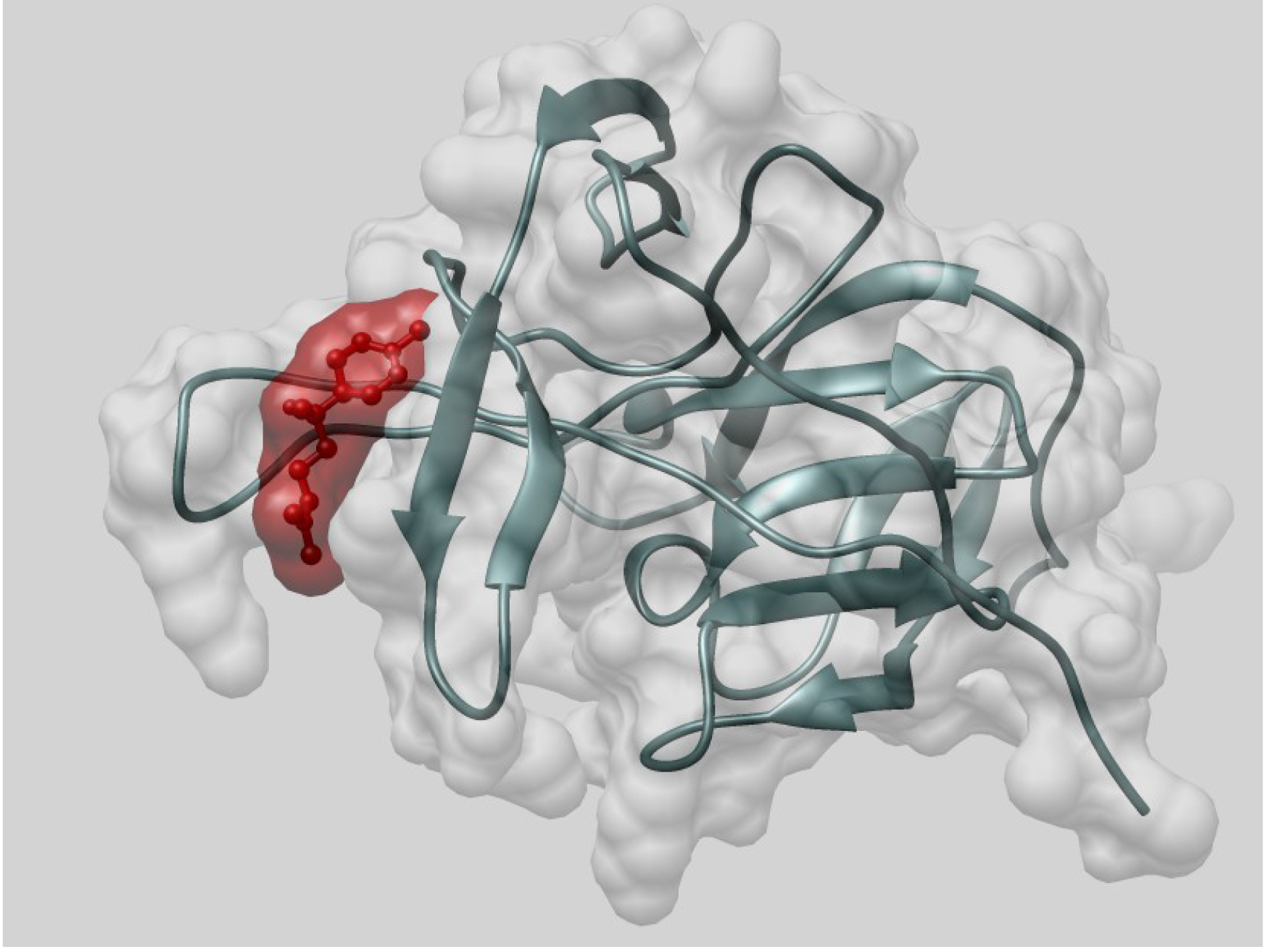
ZINC01849759 (-)-alpha-bisabolol bound to Zika virus NS3 protease(MM-PBSA Energy-18.58 kCal/mol)

**Fig. 3.**
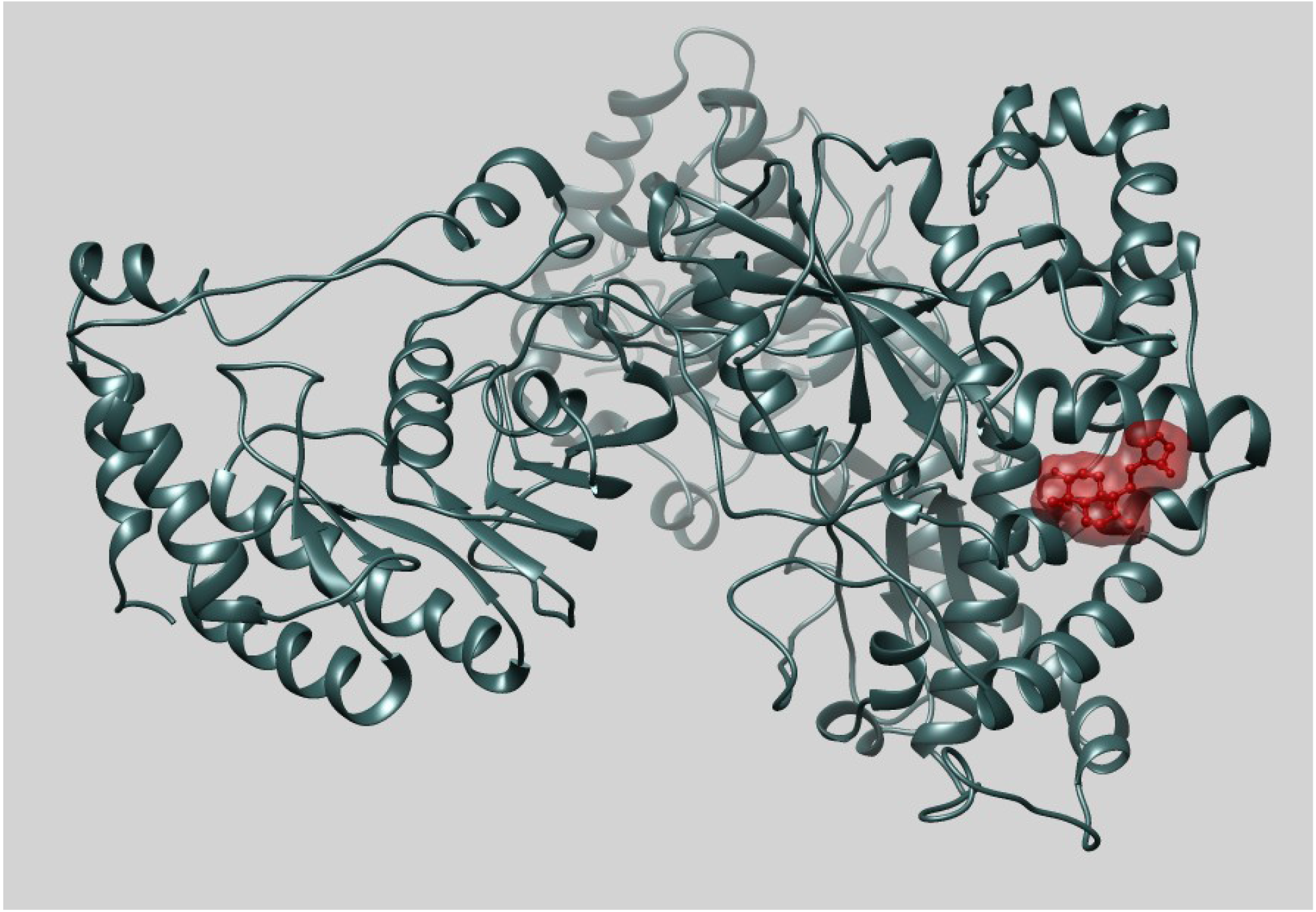
ZINC03881797 Andrographolide bound to Zika virus NS5 protein
(MM-PBSA Energy -21.14)

## Discussion

All the selected ligands demonstrate good receptor affinity in MM-PBSA experiments. The average binding affinity in MM-PBSA experiments was near to -15 kCal/mol vs. The average score of docking was on the level of -6 kCal/mol. Each of the presented ligands have some specifics in the binding. The functioning of the NS3 protease depends on the form of the cavity around the catalytic triad of the protein. The binding of eprosartan (anti-hypertension drug) changes the form of the cavity so the ferment may not perform the specific protease activity. The binding of bisabolol (phytochemical from German chamomile) to the NS3 protein competes with the binding of the NS3 protein to the NS2B cofactor, so that the protease complex may be de-activated.

According to our unpublished study, the binding site of andrographolide (phytochemical from Andrographis paniculata) in NS5 is partly formed from highly conservative residues inside each species of flaviviruses, but these residues can change between different species. The ligand which binds to this pocket can be effective for specific treatment of a particular flavivirus infection. The binding of andrographolide to the pocket of the NS5 protein has a good affinity and this substance may be the most promising antiviral agent.

Expanded work on a search for antiviral agents may include in vivo and in vitro tests of the selected substances, the search for similar ligands with higher specificity, and an expanded virtual screening by a broader base of ligands. The correct approach to an expanded virtual screening should include a clustering of ligands, selecting classes of ligands which can be suitable for inhibition of the selected proteins, and refining the search. Also, alternative vulnerable sites of the proteins can be considered, e.g. the second pocket in the NS5 polymerase or the inhibition of the NS5 methyltransferase activity. The established pipeline of analysis can be applied to a treatment of other flavivirus infections.

